# The convergence and divergence of spontaneous brain activity of humans and marmosets

**DOI:** 10.1101/2022.12.19.519301

**Authors:** Jinbo Zhang

## Abstract

The brain at rest or during periods of relative inactivity, complex dynamical patterns of activity spontaneously form over the entire brain. In this study, we plan to clarify the convergence and divergence of spontaneous brain activity across humans and marmosets. We investigated the spontaneous brain dynamics of marmosets, and compared its convergence and divergence with human neuroimaging results. We identified seven representative FC patterns (FC states) in the resting-state activity of marmosets. The most prevalent two FC patterns (VC1 and VC2) corresponds to a state of BOLD coherence of primary visual and auditory processing regions. Our results suggested that the spontaneous activity does reflect the inherent properties of the brain of marmosets.

## 1. Introduction

When our brain gets no sensory stimuli or is not engaged in a task, it is also continually active. According to research on the brain at rest or during periods of relative inactivity, complex dynamical patterns of activity spontaneously form over the entire brain (György Buzsáki, 2019). In one of the potential perspectives, spontaneous activity is similar to random noise (Shadlen & Newsome, 1998). Similar to how photons reflected from objects based on light striking the retina become available when our brain applied to the process of information, sensory representations’ content becomes accessible (Barlow, 1961).

However, another stream of idea has been suggested by mounting evidence. They believe that spontaneous activity is important for brain’s functions (see the review: Pezzulo et al., 2021). In this view, structured dynamical brain activity is related to the history of task activation and possibly serve as a prior for the execution of relevant tasks (Harmelech & Malach, 2013; Lewis et al., 2009; Petersen & Sporns, 2015). Beside, some research proposed that the spontaneous activity could be regarded the agent underlies computational models. Spontaneous fluctuations of brain activity within and across brain regions may reflect transitions between priors of the computational model (Hinton, 2007). By measuring the BOLD fMRI signal from the human brain, we have known that brain activity at rest is organized in distinct spatiotemporal patterns known as resting state networks (RSNs, see Fig. 1A). These networks are formed by groups of regions that show temporally correlated activity. According to human neuroimaging research, spontaneous activity in the hippocampus during sleep or wakeful rest mimics neuronal activation patterns for places or events encountered while awake (replays, Foster, 2017, see Fig. 1B). Furthermore, some rodent studies have suggested that at the level of single neurons within cortical and subcortical regions, spontaneous and task-evoked brain activity are similar (Pezzulo et al., 2017). These views suggested that spontaneous activity does have important physiological and cognitive significance.

**Figure 1.**
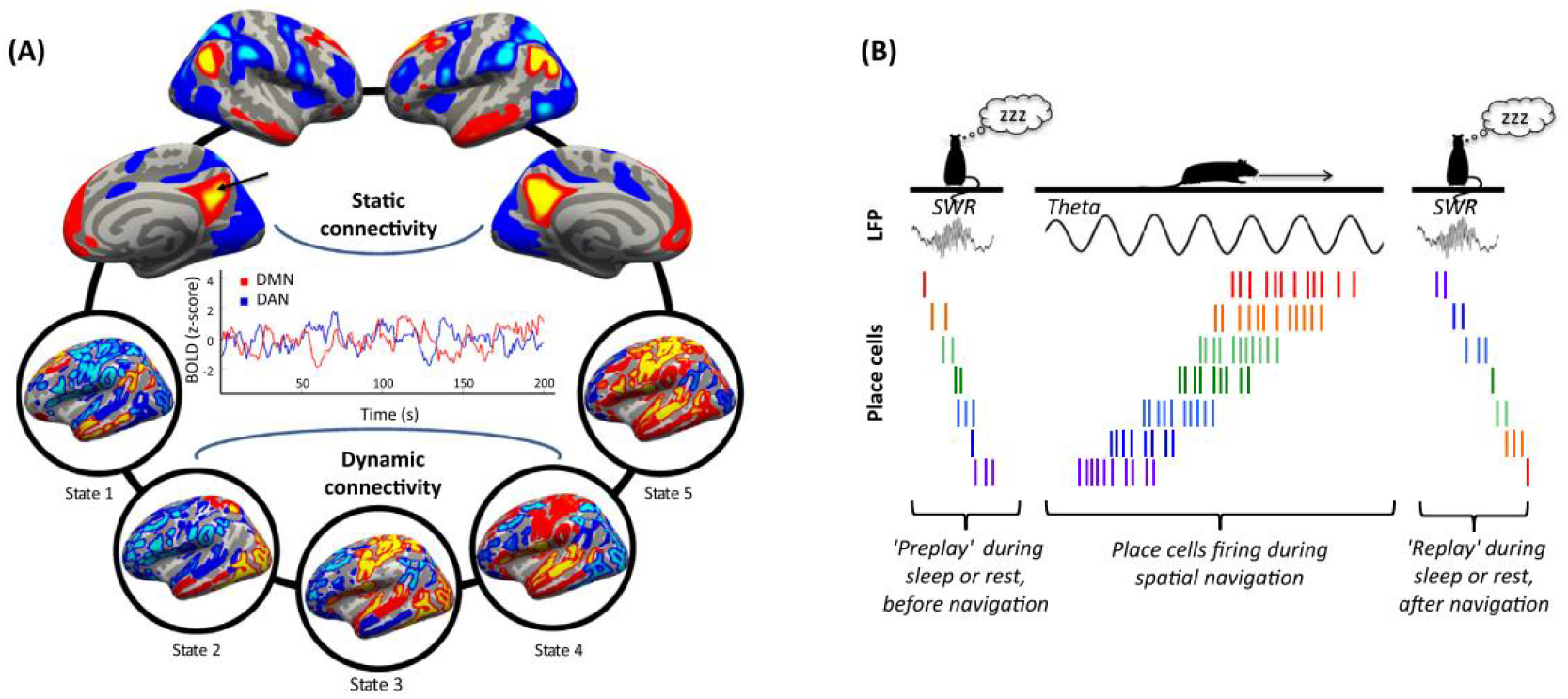
Two examples of spontaneous brain dynamics. (A) Resting state spontaneous activity measured with fMRI. Static connectivity: time-averaged temporal correlation of the blood oxygenation level-dependent (BOLD) signal between regions. Dynamic connectivity: the patterns of correlated activity across the brain as identified through a sliding window analysis. The fluctuations have a frequency of about 0.1 Hz. (B) An illustration of internally generated hippocampal sequences. The middle part of the figure shows a spatiotemporal sequence of spikes from seven hippocampal place cells, whose place fields are located in different portions of the corridor. These sequences are visible within the hippocampal theta rhythm, while the animal navigates through the corridor (Foster, 2017)

Here, I plan to clarify the convergence and divergence of spontaneous brain activity across humans and marmosets. The common marmosets (Callithrix jacchus) are a small non-human primates (NHP) species with a nearly smooth brain. Marmosets are a highly social and vocal species of New World monkeys native to South America. This feature makes marmosets very fit to be the research target of vocal communication. It can be used to reveal the behavioural and neural mechanisms of vocal communication and its disorders. A better understanding of the neurobasis of normal vocal communication will inform our understanding of how and why patients with mental (e.g., schizophrenia) or developmental (e.g., autism spectrum disorder) disorders have problems in vocal communication, thereby contributing to developing new therapeutic approaches for them. Interestingly, the resting interval of marmoset vocal communication is less studied. Considering the function importance, such as historical event replay, reconfiguration of prior of generative models, etc. it might be intesting to investigate the relationship between spontaneous brain activity and the neural basis of vocal communication.

In this study, I investigated the spontaneous brain dynamics of marmosets, and compared its convergence and divergence with human neuroimaging results. First, I used the leading eigenvector of time-resolved functional connectivity matrices based on public resting fMRI data from Marmosets Functional Connectivity described the spontaneous brain activity of marmosets (Schaeffer et al., 2022). Then different patterns of rsFC were identified with clustering analysis. This method also be used in previous study on human brain’s spontanous activity (Cabral et al., 2017). After that, the switching profiles of different brain states were investigated. The age effect was investigated by comparing the changes of brain state in young and old marmoset. In the end, the convergent and divergent features of human and marmosets’ brain activity were discussed.

## 2. Method

### 2.1 Animal

Nine adults (4 male and 5 female) marmosets (Callithrix jacchus) from the National Institutes of Health (NIH) were involved in this study, mean age of them are 54 months with a standard deviation of 22.9 months. I group them as marmosets in early adulthood (mean±standard error=31.8±10.15 months) and marmosets at the late stage of adult (age > 60 months, mean±standard error=72.8±6.1 months) based on a previous study (Sawiak et al., 2018). A non-invasive custom-formed head holder was 3D printed and affixed to the animal bed (Papoti et al., 2013). Experimental procedures were per the Canadian Council of Animal Care policy and a protocol approved by the Animal Care Committee of the National Institute of Neurological Disorders and Stroke, National Institutes of Health (NIH data). All data come from a public dataset, i.e., Marmoset Functional Connectivity.

### 2.2 Data acquisition

The NIH data were collected using a 7T 30cm horizontal bore magnet (Bruker BioSpin Corp, Billerica, MA, USA) with a custom-built 15-cm-diameter gradient coil with a maximum gradient strength of 450 mT/m (Resonance Research Corp, Billerica, MA, USA) and a custom 10-channel phased-array receive coil that conformed to the 3D printed head holder with the following parameters: TR = 2000ms, TE = 22.2 ms, flip angle = 70.4°, the field of view = 28*36mm, matrix size = 56*72, voxel size = 0.5*0.5*0.5mm, slices = 38, bandwidth = 134 kHz, GRAPPA acceleration factors: 2.

### 2.3 Data prepossessing

The fMRI data were preprocessed with the Analysis of Functional NeuroImages (AFNI) and FMRIB software library (FSL) software tools. The first ten time points were removed from magnetization to reach a steady state. The images were then despiked (AFNI’s 3dDespike), and volume was registered to the middle volume of each time series (AFNI’s 3dvolreg), and slice timing was corrected (AFNI’s 3dvolreg), and slice timing was corrected (3dTshift). Volume registration was also used to estimate head translations and rotations, which were then used as nuisance regressors and to filter movements greater than one voxel. To decrease noise, images were smoothed with a 1.5mm full-width at half-maximum (FWHM) Gaussian kernel (AFNI’s 3dmerge). Regression was used to account for variation in three-dimensional translation and rotation, as well as linear and non-linear detrending (3dDeconvolve), which also included bandpass filtering (0.01 and 0.1Hz).

### 2.4 Registration to template

The average functional image was calculated for data registered to each animal’s T2-weighted image. Anatomical images were manually skull-stripped, and this mask was applied to the functional images in anatomical space. The T2-weighted images were then nonlinearly registered to the brain atlas with 54 brain regions (Marmoset Brain Mapping, Liu et al., 2018) using Advanced Normalization Tools (ANTs).

### 2.5 Static Functional Connectivity and Dynamic Functional Connectivity

The static FC between two brain regions is measured as the Pearson correlation between their whole BOLD signals over the recording time. The FC between N = 54 brain areas is represented by an N by N matrix.

The time-resolved dynamic FC matrix (dFC) was calculated with BOLD Phase Coherence Connectivity. It is a matrix with size N by N by T, where N = 54 is the number of brain areas, T = 502 is the total number of recording frames. To calculate the phase coherence, I first estimate the phase of the BOLD signals in all areas *n*, *θ(n, t)*, using the Hilbert transform. Given the phases of the BOLD signal, the phase coherence between brain areas *n* and *p* at time *t*, *dFC(n, p, t)*, is using the following equation:

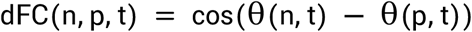

where cos() is the cosine function. Because cos(0) = 1, if two areas *n* and *p* have temporal aligned BOLD signals (i.e., they have similar phase), then dFC(n, p, t) will be close to 1. Since the phase coherence is undirected, the N by N dFC(t) matrix is symmetric. Because the matrix is symmetric across the diagonal, all relevant values may be extracted from the upper or lower triangular regions of the matrix.

### 2.6 FC Leading Eigenvector and Functional Connectivity Dynamics

The leading eigenvector *V_1_(t)* of each dFC(t) captures the dominant connectivity pattern of dFC(t) at time t, which can be reconstructed using the (N by N) outer product *V_1_V^T^_1_*.

Next, to study the evolution of the dFC over time, I compute a time-versus-time matrix representing the functional connectivity dynamics (FCD), where each entry, FCD(t_x_, t_y_), corresponds to a measure of resemblance between the dFC at times t_x_ and t_y_. The cosine similarity results in a value between −1 and 1.

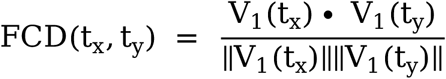

### 2.7 FC states

A discrete number of FC patterns is detected by applying clustering analysis on all the leading eigenvectors V_1_(t) across time points and subjects. I use *k*-means clustering with *k (*number of cluster) from 2 to 10, repeating each 20 times. As a result, I obtain *k* clustering solution based on the Dunn’s score.

### 2.8 Between-group comparisons

I use a Bayesian Independent Sample T-test to identify significant differences between young and old groups. This test hypothesis there is no difference in the occurrence or switching profile (dwelling time) of dFC between young and old marmoset.

### 2.9 Data and code availability statement

All data is available from Marmoset Functional Connectivity project. Codes are publicly available at http://www.github.com/Jinboasltw/dFC_marmoset

## 3. Results

### 3.1 FCD analysis

The FCD matrices reveal a characteristic pattern indicative of spontaneous switching between different recurrent FC configurations. The green squares in the diagonal of the FCD matrix represent stable FC configurations that tend to re-appear in non-contiguous time segments with sharp switches indicating a change of pattern.

### 3.2 FC states

I identify seven representative FC patterns (FC states) in the resting-state activity of all 9 marmosets (Fig. 3 and 4). More specifically, a k-means clustering algorithm was applied to the whole set of dFC leading eigenvectors and k=7 returned as the best number of FC patterns representing the data. Each of the seven cluster centroids (or states) is a vector VC, where VCVC^T^ represents an N by N connectivity pattern and VC(n) weighs the contribution of each brain area n to that pattern. FC states are ranked 1 to 7 according to their probability of occurrence, PC. I then compute the weighted sum of these VCVC^T^ matrices according to PC, and find that it strongly correlates (*r*=0.877) with the Static FC matrix averaged over all marmosets, meaning that the Static FC matrix can be fairly represented as a linear combination of only 7 eigenvectors. VC has elements with different signs, indicating that FC can be partitioned into two communities (illustrated in red and blue).

**Figure 2.**
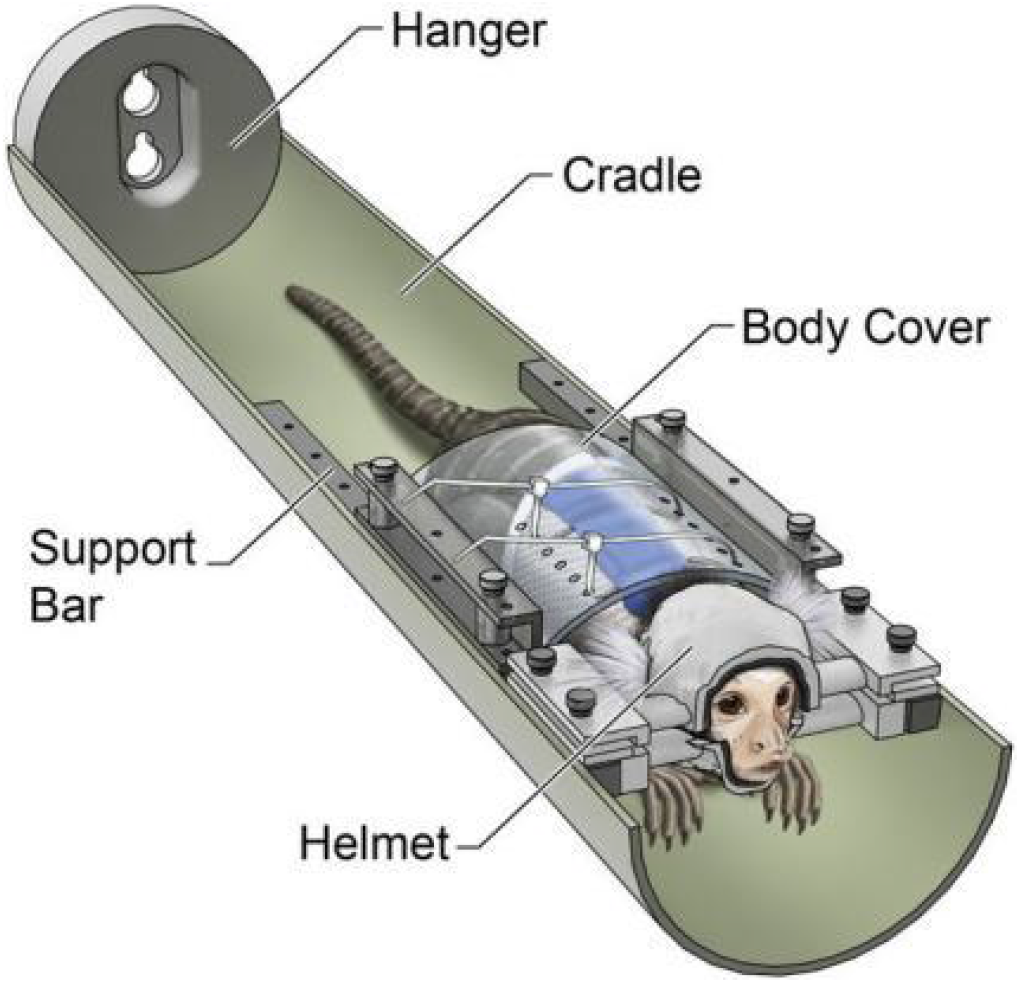
Illustration of the restraint setup used to train and image marmosets (ref Papoti et al., 2013).

**Figure 3.**
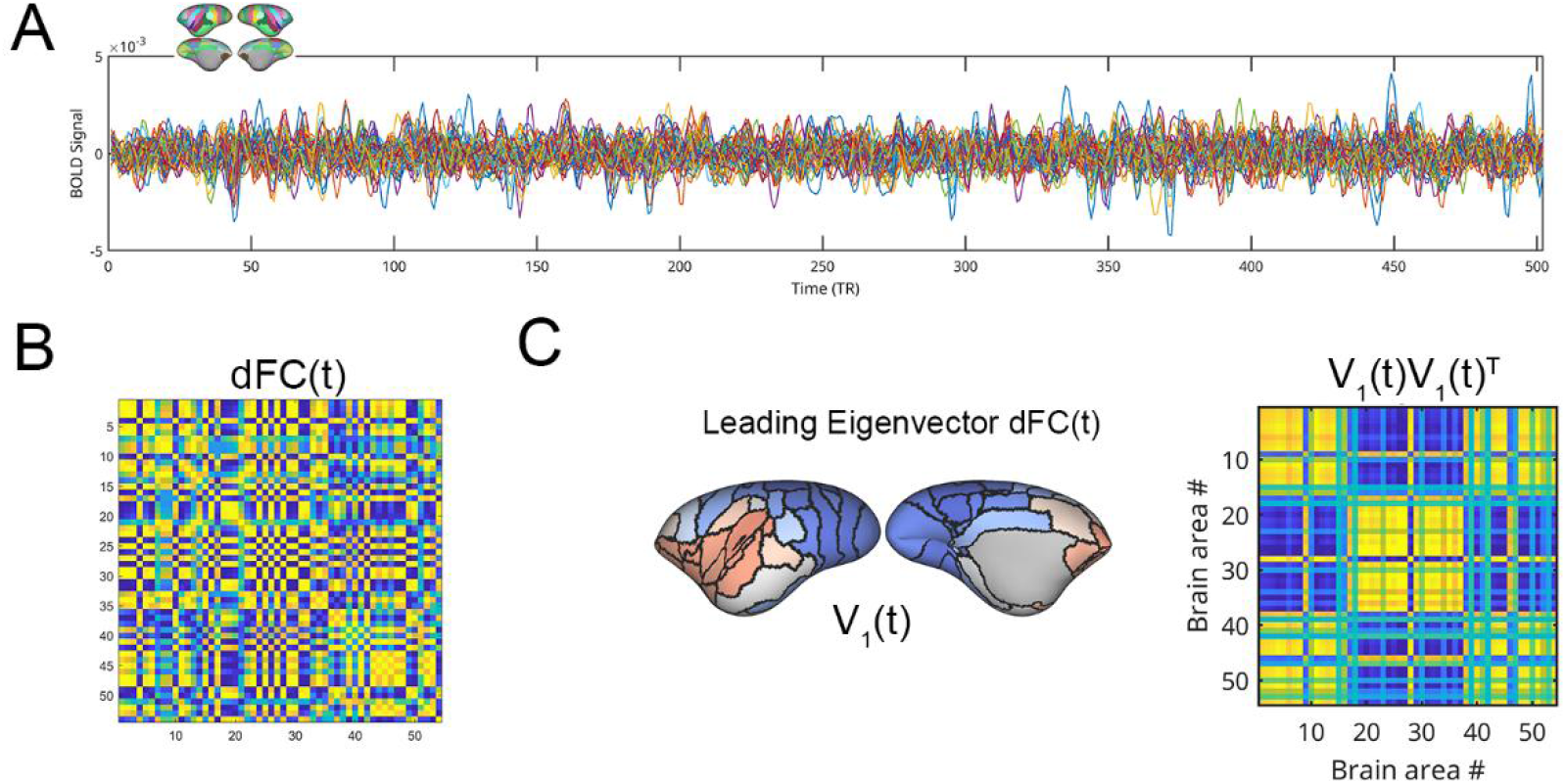
Illustration of the analysis pipeline.

**Figure 4.**
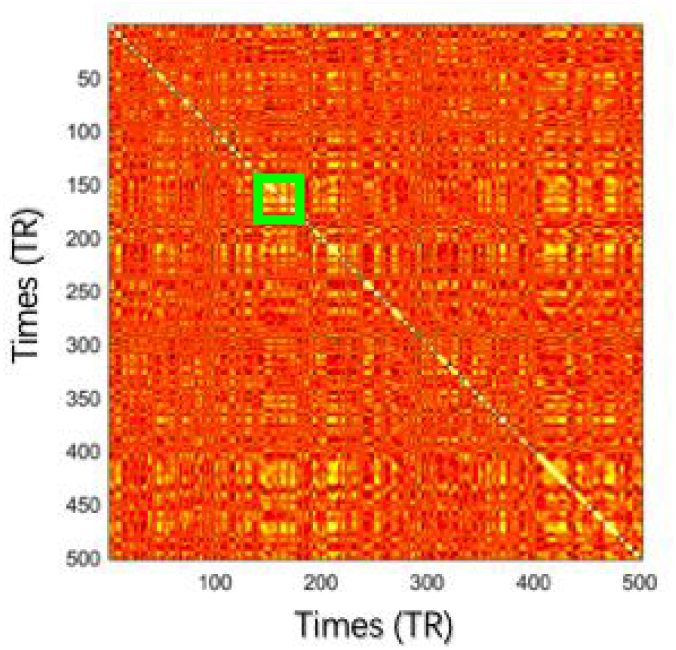
Representative result of functional connectivity dynamics (FCD).

The most prevalent two FC patterns (VC1 and VC2), which occurs more than 35% of the time, corresponds to a state of BOLD coherence of two core region: auditory belt area, superior temporal rostral area (STR), and higher-order visual association DM area (V6, at occipital lobe). In other words, during the epochs *t* when the dFC is mainly shaped by these patterns, the BOLD signals of auditory areas (Nishimura et al., 2018) and visual cortex (Rosa & Schmid, 1995) exhibit a strong coherence separately. This may suggest that auditory and visual information processing is the main functional demand for marmosets’ brains.

In the remaining FC patterns, during FC states #3–7, the BOLD phases of some subsets of brain areas misalign from the rest of the network and temporarily align together. In FC pattern #3, I find two communities. One is consisting of the somatosensory cortex (S1), and another is formed based on areas from the medial prefrontal cortex (A32). In FC pattern #4, the dorsolateral prefrontal cortex (DLPFC), medial prefrontal cortex (MPFC), and medial superior temporal area (MST) form a functional network independent from the rest of the brain. Notably, it covered typical regions from the Default Mode Network. FC patterns #5 and #6 represent a decoupling of frontal from sensory-motor-related cortices. Finally, FC pattern #7 is characterized a functional network of fundal superior temporal area (FST). It plays an important role in motion perception and direction selectivity (Hinton, 2007).

### 3.3 Between-group differences in FC patterns

To investigate the relationship with age, I compare the FC dwell time between young and old adult marmosets in each state. During the FC state#4, the resting-state FC in young marmosets is more stable in the sense that FC states last longer (mean±standard error=2.52±0.05 TRs, Bayes factor, BF_10_=24.4 > 3) whereas FC states last shorter in older adult marmosets (2.074±0.07 TRs) (Fig. 6). No other significant differences were found (BF_10_ < 3).

**Figure 5.**
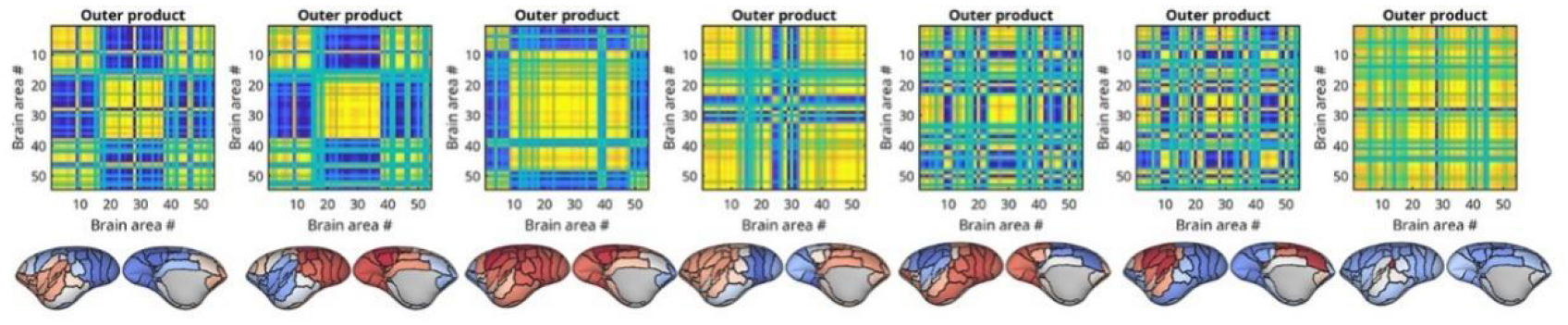
Brain state (left to right: state#01 to state#07) and corresponding coherence pattern (output product of VCVC^T^).

**Figure 6.**
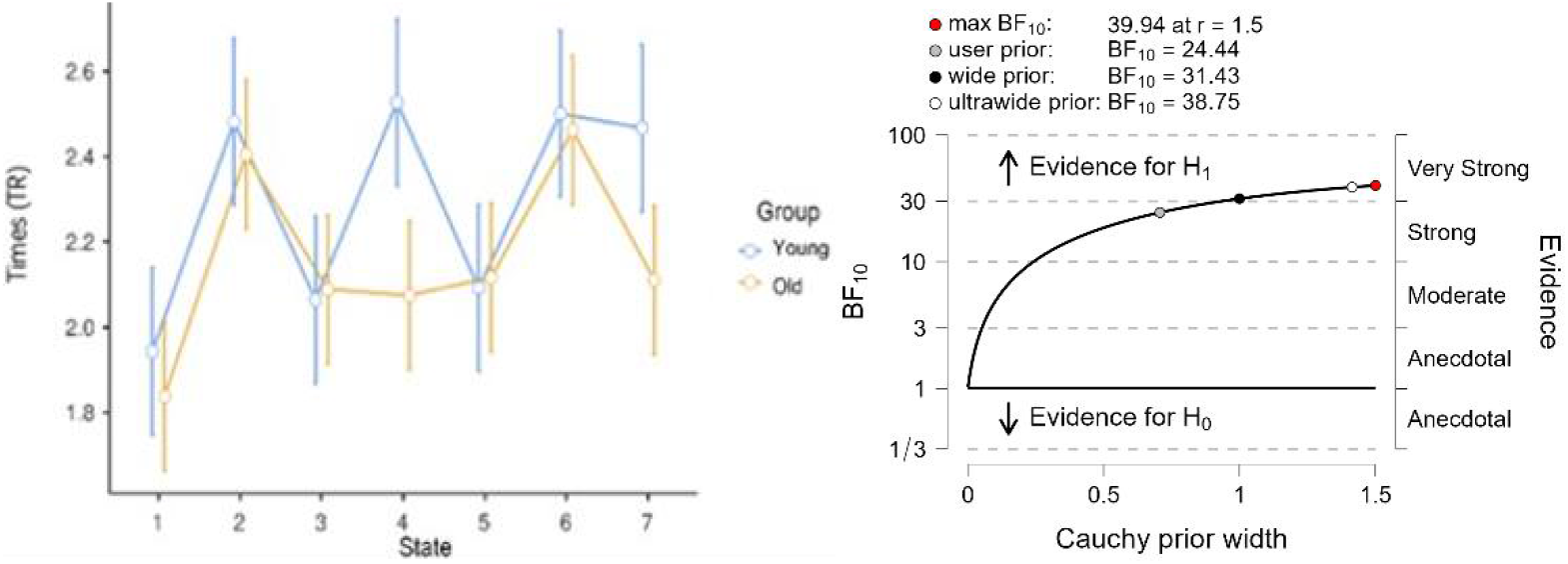
Comparing of dwell time among young and old marmosets at each state of brain.

## 4 Discussion and Conclusions

In the present work, I find that the switching behaviour of resting-state FC in marmosets relates to their marmoset’s age or the stage of adulthood. These results are in line with the idea that brain activity relates to high-order processes deeply involved in individual development and ageing (Sporns et al., 2004).

Here, I find a clear relationship between the prevalence of FC state #4 and age, pointing to a potential role of frontal BOLD coherence in cognitive processing, which lasts shorter and occurs with less probability in old marmosets. The lower occurrence of this state in old marmosets, in line with previous findings of human studies where decrease in BOLD functional connectivity of frontal-parietal cortical in elder people (typical ageing and atypical ageing, Chen et al., 2017; Gu et al., 2020).

Compared with human spontaneous activities, marmoset has more types of spontaneous activities, more frequent switching and shorter duration. But even so, it was found that the state of spontaneous activity changed with development and ageing. This means that spontaneous activity does reflect the inherent properties of the brain of marmosets.

There are some limitations in this study. First of all, compared with human research, it is very convenient to test task-state fMRI, but for animal fMRI, it is difficult to carry out experimental research. Therefore, it is difficult to explain the clear relationship between these spontaneous activity states and cognition. In addition, the time resolution of magnetic resonance imaging is lower than that of EEG, which means that it is difficult to capture spontaneous activity with a very short duration (MilliTR). This study found that marmoset’s brain state is more and shorter in comparing with the spontaneous state of humans. Therefore, in the future, the spontaneous brain activity of marmoset can be described more precisely with the high time resolution of electrophysiological records. Besides, considering the function importance, such as historical event replay, reconfiguration of prior of generative models, etc, it might be also important to investigate the relationship between spontaneous brain activity by combining it with study of replay.

